# Impact of Inactivation Methods on Biosafety and Antigen Reactivity of *Brucella melitensis* from the Perspective of Astral-DIA Proteomics Based on Antibody Immunoprecipitation Mass Spectrometry

**DOI:** 10.1101/2025.07.28.667132

**Authors:** Yijian Liu, Jiazhen Ge, Guodong Song, Pengcheng Gao, Mengzhu Qi, Wenhao Wang, Yingying Xie, Ziqing Wang, Renge Li, Yuefeng Chu, Fuying Zheng

## Abstract

Effective *Brucella* inactivation is imperative for safe vaccine development, diagnostics, and sample handling, particularly in resource-limited regions lacking high-level containment facilities. This study investigated inactivation methods using the Rev.1 vaccine strain and three *Brucella melitensis* field isolates from Gansu (GS-XG, GS-SN, GS-MQ). Heat (80°C/95°C, 10-20 min) and formaldehyde (0.4%/0.6%, 48-72 h) inactivation were evaluated for biosafety and antigenicity. Rev.1 was completely inactivated by 80°C/95°C for 20 min or 0.6% formaldehyde for 48-72 h. Superior antigenicity, compared to phenol inactivation, was confirmed by ELISA and Western Blot. However, field isolates demonstrated greater resistance, surviving 80°C for 20 min and 0.4% formaldehyde for 72 h, necessitating stricter conditions (95°C, 20 min; 0.6% formaldehyde, 72 h) for their complete inactivation. Astral-DIA proteomics analyzed approximately 60% of the proteome (∼2000/3300 proteins), revealing 256, 311, and 318 differentially expressed proteins between 80°C/95°C heat, 48-h/72-h formaldehyde, and heat/formaldehyde methods, respectively. Gene Ontology and KEGG analyses indicated heat inactivation upregulated cellular structure proteins but downregulated metabolic pathways, with 95°C potentially damaging conformational epitopes. Formaldehyde treatment (48 h) stabilized soluble antigens, preserving ribosomal and regulatory protein epitopes, while 72-h treatment induced organelle disintegration. Protein-Protein Interaction networks suggested heat inactivation enhanced immunogenicity of membrane and stress proteins, suitable for targeted studies, whereas formaldehyde preserved broader epitopes, beneficial for whole-cell vaccines and multi-epitope screening. Inactivation methods must be tailored to specific strain characteristics and applications. Astral-DIA provides molecular insights into antigenicity loss, guiding future research on protein functions and epitope dynamics for precise brucellosis control.

## Introduction

*Brucella*, a facultative anaerobic intracellular bacterium, encompasses six recognized classical species: *Brucella. melitensis*, *Brucella. abortus*, *Brucella. suis*, *Brucella. neotomae*, *Brucella. ovis*, and *Brucella. canis*. Among these, *B. melitensis*, *B. abortus*, and *B. suis* are capable of infecting both their natural hosts and humans, collectively causing the zoonotic disease known as brucellosis [1]. Approximately 500,000 individuals are affected annually, posing a significant public health challenge, particularly in developing countries where the disease is prevalent in both animal and human populations [2]. In animals, infection is primarily acquired through the ingestion of contaminated food or water, often tainted by infected tissues such as aborted fetuses or placental membranes. Human infections typically result from direct contact with the blood or tissues of infected animals, or from the consumption of contaminated dairy products, such as unpasteurized milk and cheese [3]. Furthermore, aerosolized pathogens can lead to respiratory tract infections, and improper laboratory practices are known to precipitate occupational exposures [4]. This multi-species, multi-route transmission characteristic contributes to a cyclical transmission chain within livestock and endemic areas, thereby increasing the complexity of control and prevention efforts. Economic losses in animals are substantial, primarily due to infertility and abortions caused by brucellosis [5]. In humans, the acute phase of the disease is characterized by undulating fevers, followed by a chronic phase that can affect most organs, manifesting in symptoms such as arthritis, orchitis, hepatitis, meningoencephalitis, and endocarditis [6]. Despite the implementation of livestock screening and vaccination strategies for brucellosis control in some developing countries in the Middle East, Asia, Africa, and South America, the disease has yet to be effectively controlled or eradicated [7].

The inactivation of *Brucella* plays a pivotal role in various experimental contexts, including vaccine development, diagnostic reagent preparation, and the collection of veterinary samples at the primary care level. In vaccinology, inactivation techniques are employed to disrupt the pathogen’s replicative capacity through heat, chemical agents, or genetic engineering, while concurrently preserving its immunogenicity [8]. For instance, the genetically attenuated *Brucella abortus* S19 strain, which is inactivated by gene deletion, can be used to produce safe vaccines. The characteristic lipopolysaccharide (LPS) antigen deletion in this strain prevents the production of interfering antibodies in immunized animals, thereby facilitating the differentiation between natural infection and vaccination through serological testing [4]. In diagnostic reagents, LPS antigens extracted from inactivated bacteria serve as a core component for colloidal gold immunoassay kits. Their stable antigenic epitopes ensure high sensitivity and specificity in detection [9]. Furthermore, novel inactivation technologies, such as *Brucella* bacterial ghosts combined with mRNA delivery systems, have been developed to enhance the protective efficacy of inactivated vaccines, addressing the limitations of insufficient immunogenicity associated with traditional inactivated vaccines [10]. During the collection of veterinary samples at a primary level, inactivation procedures are crucial for mitigating biosafety risks associated with *Brucella* during sample transportation and subsequent testing. This prevents potential infection of personnel and sample contamination due to residual live bacteria, thereby ensuring the accuracy and safety of downstream laboratory analyses. Current research trends indicate that the synergistic application of inactivated vaccines with molecular marker technologies (e.g., VirB12 gene deletion) can optimize immune protection rates (e.g., the A19-ΔVirB12 strain has demonstrated over 80% protection against challenge in calves), while also supporting the achievement of eradication goals in endemic areas [11].

The selection of inactivation methods for *Brucella*, a pathogen posing a significant threat to both human and animal health, is critical. This choice directly impacts biosafety, the efficacy of downstream applications, and proteomic alterations. Existing research indicates that methods such as heat inactivation (80°C for 5-10 minutes) and formaldehyde inactivation (0.6% formaldehyde for 72 hours) are effective in rendering the pathogen non-viable while largely preserving its immunogenicity. These methods have been successfully employed in the preparation of safe vaccines and diagnostic reagents [12]. However, formaldehyde inactivation necessitates precise concentration control; insufficient concentrations may lead to incomplete inactivation, while higher concentrations exhibit variable efficacy across different *Brucella* strains [13]. Although commonly used disinfectants can rapidly eliminate the pathogen, differential sensitivities among *Brucella* species have been observed. Furthermore, physical, chemical, and emerging biotechnological inactivation methods not only present safety concerns due to potential incomplete inactivation or chemical residues but can also induce changes in the bacterial proteome, thereby affecting antigenic reactivity and product preparation outcomes [14]. In laboratory settings, *Brucella* is classified as a Biosafety Level 3 (BSL-3) pathogen. Should inactivation be incomplete, viable bacteria can lead to researcher infections through routes such as aerosol transmission or sharps injuries [15]. Consequently, a systematic comparison of different inactivation methods, particularly regarding their effects on biosafety and proteomic profiles, is essential. Identifying safe and effective inactivation protocols is crucial for future research, significantly contributing to the improvement of brucellosis control and prevention strategies, and ultimately safeguarding both human and animal health.

This study specifically investigates the impact of various *Brucella* inactivation methods on both biosafety and antigenic reactivity. The primary objectives are to identify the inactivation approach that maximally ensures biosafety, thereby mitigating the risk of infection arising from incomplete inactivation, and to determine the method that optimally preserves antigenic reactivity. The latter is crucial for eliciting robust protective immunity in vaccine development and for achieving accurate detection in diagnostic reagent preparation. Through this research, scientifically sound references are intended to be provided, enhancing the safety and applicability of *Brucella*-related procedures, including vaccine production and laboratory testing.

## 1 Materials and Methods

### 1.1 Strains and materials

The *Brucella. melitensis* strains utilized in this study were as follows: Rev.1, which was preserved at the Lanzhou Veterinary Research Institute’s Strain Collection Center, and three field isolates: GS-XG, GS-SN, and GS-MQ. These field isolates were obtained from naturally infected animals in Xigu, Sunan, and Minqin, Gansu Province, China respectively.

Tryptic Soy Broth (TSB) and Tryptic Soy Agar (TSA) (BD, America). Reagents including 10% formaldehyde, phenol, H_2_O_2_ solution, and PBS powder (Coolaber, China). Skim milk powder (Beyotime, China). HRP-conjugated rabbit anti-goat polyclonal antibody (Sigma, America).

### 1.2 Preliminary screening of strain inactivation conditions

Preliminary inactivation screening experiments were conducted using the methods outlined in Table 1. Rev.1 bacterial strains, preserved in glycerol, were initially inoculated onto TSA medium and incubated at 37°C for five days. Subsequently, a single colony was selected and transferred to TSB medium for cultivation at 37°C with shaking at 180 r/min for 72 hours. Following this, the culture was streaked onto TSA medium and incubated at 37°C for an additional five days, after which the number of viable bacteria in the broth was enumerated. The bacterial suspension was then diluted with TSB to a concentration of 1 × 10^11 CFU/mL and stored for further use.

To investigate the effect of different inactivation methods on the antigenic reactivity of bacterial strains, initial screening was performed using ELISA to identify inactivated bacterial solutions that exhibited good reactivity with serum, based on the absence of colony growth from the previously obtained results. Briefly, the bacterial solution was centrifuged and the pellet was resuspended in PBS. Phenylmethanesulfonyl fluoride (PMSF) was then added, and the mixture was thoroughly agitated on ice. Complete bacterial lysis was achieved through sonication at 0°C for 20 minutes (using a 6mm probe at 39% maximum power, with pulses of 2 seconds on and 3 seconds off). Subsequently, the solution was centrifuged at 12,000 g for 30 minutes at 4°C to remove insoluble material, and the supernatant, containing the total bacterial protein solution, was collected. This solution was then used to coat 96-well plates. Following an overnight coating period, plates were washed three times. Blocking was performed at 37°C for 1 hour. After three washes, positive and negative sera were added as primary antibodies and incubated at 37°C for 1 hour. Plates were washed three times, and HRP-conjugated rabbit anti-goat polyclonal antibody (1:10,000 dilution) was added and incubated at 37°C for 1 hour. After a final three washes, chromogenic substrate was added in the dark. Color development was allowed to proceed for 20 minutes prior to the addition of stop solution. Subsequently, optical density (OD) values were measured at a wavelength of 450 nm using an enzyme-linked immunosorbent assay (ELISA) workstation (Tecan, Switzerland).

### 1.3 Thermal and formaldehyde inactivation methods were applied to various clinically isolated bacterial strains

Field isolates 2 and 3 were subjected to inactivation under identical conditions using two primary methods: heat inactivation and formaldehyde inactivation. The specific methodologies are detailed below.

#### Heat Inactivation

Four 100 mL aliquots of cultured bacterial suspension were prepared for inactivation. These were then subjected to water bath treatment at 80°C and 95°C for 10 minutes and 20 minutes, respectively. Following inactivation, an appropriate volume of each treated bacterial suspension was inoculated onto TSA agar plates and incubated at 37°C for one week to assess for colony growth. The remaining inactivated bacterial suspensions were centrifuged at 8000 rpm, the supernatant was discarded, and the pellets were stored at 4°C.

#### Formaldehyde (HOCO) Inactivation

Four 100 mL aliquots of cultured bacterial suspension were prepared for inactivation. To each, 10% formaldehyde solution was added to achieve final concentrations of 0.4% and 0.6%, respectively. The mixtures were then incubated at 37°C. For the 0.4% formaldehyde concentration, an appropriate volume of inactivated bacterial suspension was sampled at 72 hours. For the 0.6% formaldehyde concentration, samples were taken at 48 hours and 72 hours. These samples were then inoculated onto TSA agar plates and incubated at 37°C for one week to observe for colony growth, thereby confirming inactivation. The remaining inactivated bacterial suspensions were centrifuged at 8000 rpm, the supernatant was discarded, and the pellets were stored at 4°C.

### 1.4 Astral-DIA Immunoprecipitation Proteomic

#### 1.4.1 Inactivation procedure

GS-XG strain was selected for these experiments, its moderate virulence having been established in unpublished studies. Inactivation methods were optimized through a time-gradient approach, encompassing treatments at 80°C for 20 minutes, 95°C for 20 minutes, 0.6% formaldehyde for 48 hours, and 0.6% formaldehyde for 72 hours. Prior to subsequent immunoprecipitation (IP) and mass spectrometry analyses, all bacterial strains underwent a 7-day biosafety verification period.

#### 1.4.2 Immunoprecipitation and Astral-DIA Proteomics

Four sets of inactivated samples (80°C for 20 min, 95°C for 20 min, 0.6% formaldehyde for 48 h, and 0.6% formaldehyde for 72 h) were processed. Whole-cell protein extracts, obtained by sonication, were subjected to immunoprecipitation (IP) using Protein A/G magnetic beads (APEXBIO, USA). Chromatographic separation was performed on a nano-flow VanquishNeo system (ThermoScientific, USA). Subsequently, samples eluted from the nano-scale high-performance liquid chromatography were analyzed by DIA (data-independent acquisition) mass spectrometry using an Orbitrap Astral platform (ThermoScientific, USA) (Petrosius, et al., 2023). Detection parameters were set as follows: MS1 was acquired from 380–980 m/z with a resolution of 240,000 (at 200 m/z) and a maximum injection time of 5 ms; MS2 acquisition involved 299 windows, an isolation window of 2 m/z, HCD collision energy of 25 eV, and a maximum injection time of 3 ms. DIA data were processed using DIA-NN software (Demichev, et al., 2019). The software parameters were configured as follows: trypsin was designated as the enzyme, with a maximum of one missed cleavage site. Carbamidomethyl (C) was set as a fixed modification, while Oxidation (M) and Acetyl (Protein N-term) were defined as dynamic modifications. Identified proteins were required to meet a false discovery rate (FDR) of <1% for acceptance.

### 1.5 Bioinformatics analysis

#### 1.5.1 Protein annotation and classification

Gene Ontology annotations for proteins were generated using EggNOG-mapper (v2.0) against the EggNOG 5.0 database, categorized by cellular component, molecular function, and biological process [16]. Protein conserved domains were identified through the PfamScan tool and the Pfam database [17]. Pathway annotations were performed by selecting the highest-scoring results from BLASTP (e-value ≤ 1e⁻⁴) alignments against the KEGG database. Subcellular localization was determined using PSORTb (v3.0), analyzed specifically for Gram-negative bacteria [18]. Homologous protein clusters were categorized based on the EggNOG database.

#### 1.5.2 Functional enrichment analysis

Enrichment analyses for Gene Ontology (GO), KEGG pathways, and protein domains were performed using Fisher’s exact test. A significance threshold of P < 0.05 was applied [19]. Domain enrichment was specifically based on the Pfam database.

#### 1.5.3 Protein–protein interaction (PPI) networks analysis

Differential proteins were mapped against the STRING (v11.0) database using the wemol supercomputing platform (https://wemol.wecomput), with a confidence score exceeding 0.7 [20]. Interaction networks were subsequently constructed utilizing the R package visNetwork (https://github.com/datastorm-open/visNetwork).

### 1.6. Statistical analysis

All experiments were independently repeated at least three times, and each sample was analyzed three times. Bacterial count was conducted using the calculation of average (Mean) and standard deviation (Standard Deviation, SD).

## 2 Result

### 2.1 Formalin- and heat-inactivated *B. melitensis* Rev.1 exhibited commendable safety and antigenicity

Plate counts for various inactivation methods are presented in Table 2. It was observed that the Rev.1 vaccine strain consistently yielded no colony formation, regardless of inactivation conditions. These conditions included water bath heating at 80°C for 10 or 20 minutes, 95°C for 20 minutes, treatment with 0.4% formalin for 72 hours, 0.6% formalin for 48 or 72 hours, or inactivation with 2% phenol for 10 minutes or 5% phenol for 10 minutes.

The antigenicity of the inactivated *Brucella* Rev.1 strain was assessed by ELISA (Figure 1A). It was determined that both formalin- and heat-inactivated preparations exhibited superior antigenicity compared to phenol-inactivated preparations. This finding was further corroborated by subsequent Western blot analyses (Figure 1B and C).

**Figure 1:**
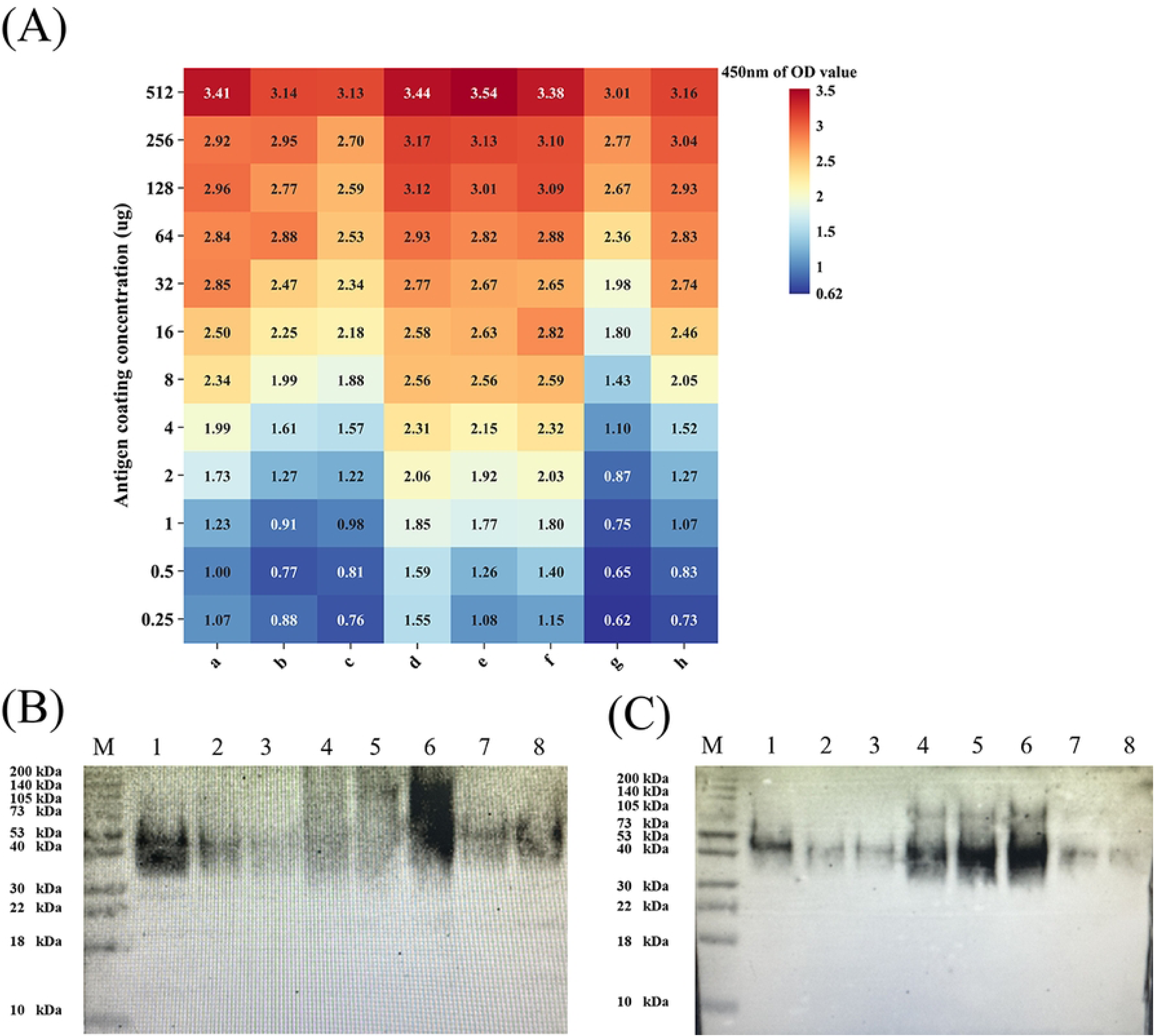
(A) Effect of eight inactivation conditions on protein antigenicity, as measured by OD450nm. Conditions are represented as follows: a: 80°C for 20 min; b: 95°C for 20 min; c: CHCA; d: 0.4% HCHO for 72 h; e: 0.6% HCHO for 48 h; f: 0.6% HCHO for 72 h; g: 2% phenol for 10 min; h: 5% phenol for 10 min. (B) Western Blot validation o f the reactivity of proteins with infected serum under various inactivation conditions. Lane assignments are as follows: Line M: protein marker; Line1: 80°C for 20 min; Line2: 95° C for 20 min; Line3: CHCA; Line4: 0.4% HCHO for 72 h; Line5: 0.6% HCHO for 48 h; Line6: 0.6% HCHO for 72 h; Line7: 2% phenol for 10 min; Line8: 5% phenol for 10 min. (C) Western Blot validation of the reactivity of proteins with immune serum under v arious inactivation conditions. Sample assignments for each lane are identical to those in p anel (B).

### 2.2 Biosafety concerns arose when the conditions effective for inactivating the Rev.1 vaccine strain were applied to clinical isolates

Given biosafety considerations and antibody profiling, heat inactivation and formalin inactivation were selected for subsequent experiments. Both heat and formalin treatments demonstrated complete inactivation of all tested isolates, and their antigenicity retention, as evidenced by ELISA and Western blot, was significantly superior to methods employing phenol and similar agents. These two methods exhibited a notable advantage in balancing inactivation efficacy with antigenic integrity, thus providing an ideal model for subsequent proteomic analysis.

However, when these methods were applied to clinical isolates, perfect inactivation was not achieved for all strains (Table 3). Plate counts revealed colony growth for isolate 2 and isolate 3 following inactivation at 80°C. Conversely, no colony growth was observed for isolate 2 and isolate 3 after high-temperature inactivation at 95°C for both 10 and 20 minutes, indicating complete inactivation. Plate counts further indicated that isolate 2 and isolate 3 remained viable after treatment with 0.4% formalin for 72 hours. Complete inactivation of isolate 2 and isolate 3 was achieved with 0.6% formalin after both 48 and 72 hours, as no colonies were observed on solid media.

### 2.3 Identification and Distribution Characteristics of Differentially Expressed *Brucella* Proteins under Various Inactivation Conditions

To balance biosafety and antigenicity, *Brucella* samples inactivated under four distinct conditions—80°C for 20 min, 95°C for 20 min, 0.6% formaldehyde for 48 h, and 0.6% formaldehyde for 72 h—were selected for subsequent IP experiments. Mass spectrometry analysis was performed using an Orbitrap Astral platform.

High signal intensities were observed in the mass-to-charge ratio (m/z) range of 500 to 1500 for samples treated with 80°C for 20 min, 95°C for 20 min, 0.6% formaldehyde for 48 h, and 0.6% formaldehyde for 72 h (Figure 2A, B, C, D). Specifically, the 80°C/20 min group exhibited characteristic strong peaks at m/z 536.17 and 610.19 (Figure 2A). The 95°C/20 min group showed significant signal clusters at m/z 421.76 and 537.17 (Figure 2B). Dominant ion peaks were observed at m/z 445.12 and 538.16 in the 0.6% formaldehyde/48 h group (Figure 2C), while high-intensity signal regions were present at m/z 528.34 and 738.04 in the 0.6% formaldehyde/72 h group (Figure 2D). For quantitative purposes, only proteins with report intensities from at least two of the three biological replicates were included. A total of 2054, 2053, 2041, and 2015 quantifiable proteins were identified in the 80°C/20 min, 95°C/20 min, 0.6% formaldehyde/48 h, and 0.6% formaldehyde/72 h groups, respectively (Figure 2E). Overall, 1750 proteins were commonly detected across all four sample groups (Figure 2E). Furthermore, 1788 proteins were shared between the 80°C/20 min and 95°C/20 min groups, and 1763 proteins were shared between the 0.6% formaldehyde/48 h and 0.6% formaldehyde/72 h groups.

**Figure 2:**
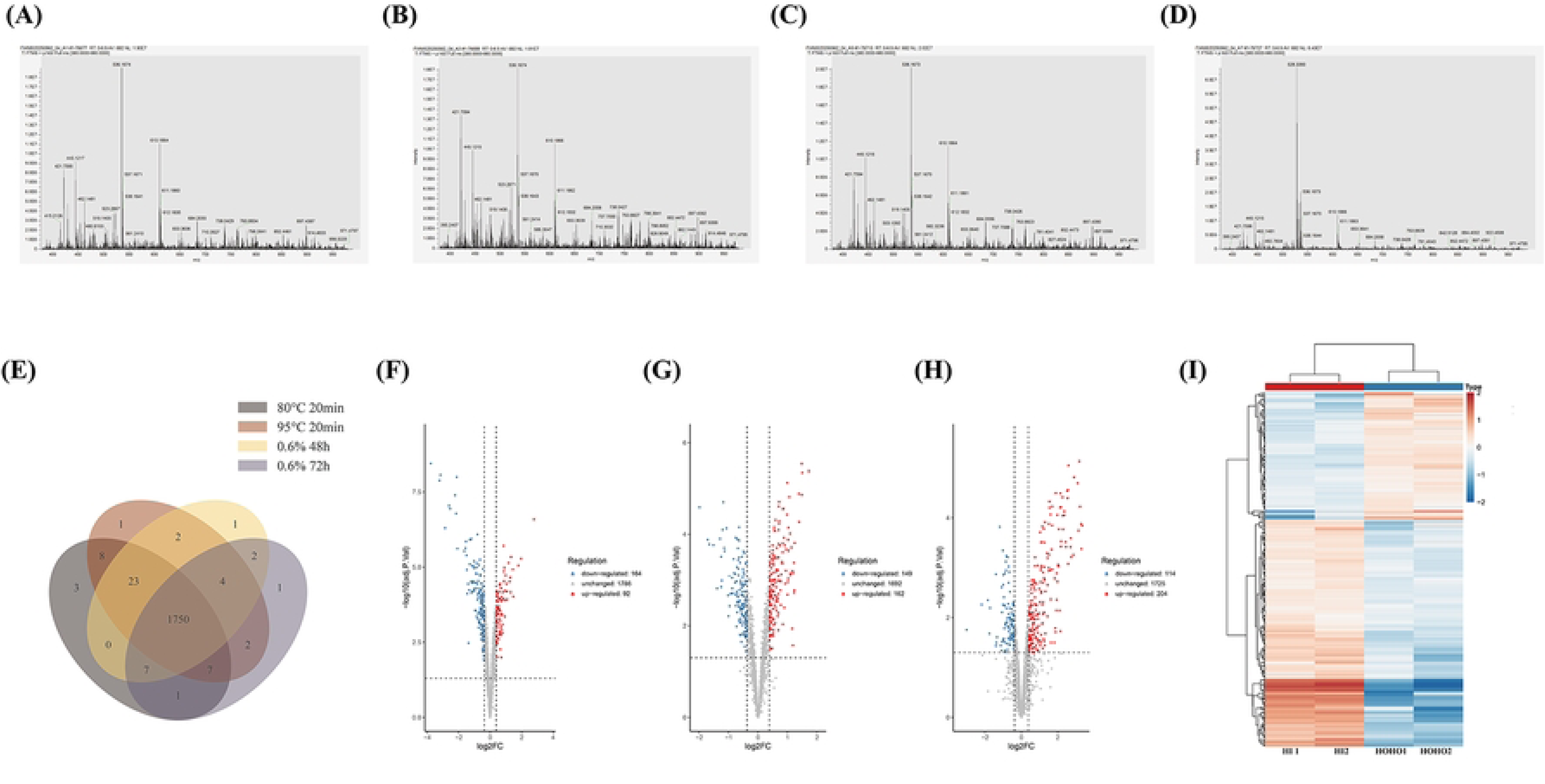
(A) MALDI-TOF mass spectrum of the sample inactivated at 80°C for 20 min. (B) MALDI-TOF mass spectrum of the sample inactivated at 95°C for 20 min. (C) MA LDI-TOF mass spectrum of the sample inactivated with 0.6% HCHO for 48 h. (D) MAL DI-TOF mass spectrum of the sample inactivated with 0.6% HCHO for 72 h. (E) Overlap of protein identifications across four distinct inactivation methods. (F) Volcano plot illustr ating differentially expressed proteins between the 80°C/20 min and 95°C/20 min comparat ive groups. (G) Volcano plot illustrating differentially expressed proteins between the 0.6% HCHO/48 h and 0.6% HCHO/72 h comparative groups. (H) Volcano plot illustrating diff erentially expressed proteins between the heat-inactivated and formaldehyde-inactivated grou ps. (I) Heatmap illustrating differentially expressed proteins between the heat-inactivated an d formaldehyde-inactivated groups.

Through Astral-DIA antibody IP proteomic analysis, 256 differentially expressed proteins were identified between the 80°C/20 min heat inactivation group and the 95°C/20 min heat inactivation group. Among these, 92 proteins exhibited significantly higher expression in the 80°C/20 min group compared to the 95°C/20 min group, while 164 proteins showed significantly lower expression (Figure 2F). Differential analysis between the 0.6% formaldehyde/48 h and 0.6% formaldehyde/72 h groups revealed 311 differentially expressed proteins. Of these, 162 proteins were found to be significantly more abundant in the 0.6% formaldehyde/48 h group than in the 0.6% formaldehyde/72 h group, and 149 proteins were significantly less abundant (Figure 2G).

A total of 318 differentially expressed proteins were identified between the heat-inactivated (20 min) and 0.6% formaldehyde-inactivated groups. Among these, 204 proteins were significantly more abundant in the heat-inactivated (20 min) group compared to the 0.6% formaldehyde-inactivated group, while 114 proteins were significantly less abundant (Figure 2H). The heatmap distribution indicated clear distinctions between the heat-inactivated (20 min) and 0.6% formaldehyde-inactivated groups, with consistent coloring within each subgroup, demonstrating good reproducibility (Figure 2I). Clustering analysis revealed that most proteins upregulated by heat inactivation were downregulated by formaldehyde inactivation, and vice versa (Figure 2I). All the relevant information regarding the differences in expressed proteins can be found in the supplementary materials.

### 2.4 Functional Classification Based on GO/KOG/KEGG Enrichment

GO functional classification of differentially expressed proteins revealed that proteins with higher expression in the 80°C/20 min group relative to the 95°C/20 min group were significantly enriched in catalytic activity (6 genes), response to stimulus (3 genes), metabolic processes (5 genes), and cellular processes (5 genes). Conversely, proteins with lower expression in the 80°C/20 min group compared to the 95°C/20 min group were significantly enriched in catalytic activity (13 genes), cellular anatomical entities (19 genes), metabolic processes (15 genes), and cellular processes (15 genes) (Figure 3A).

**Figure 3:**
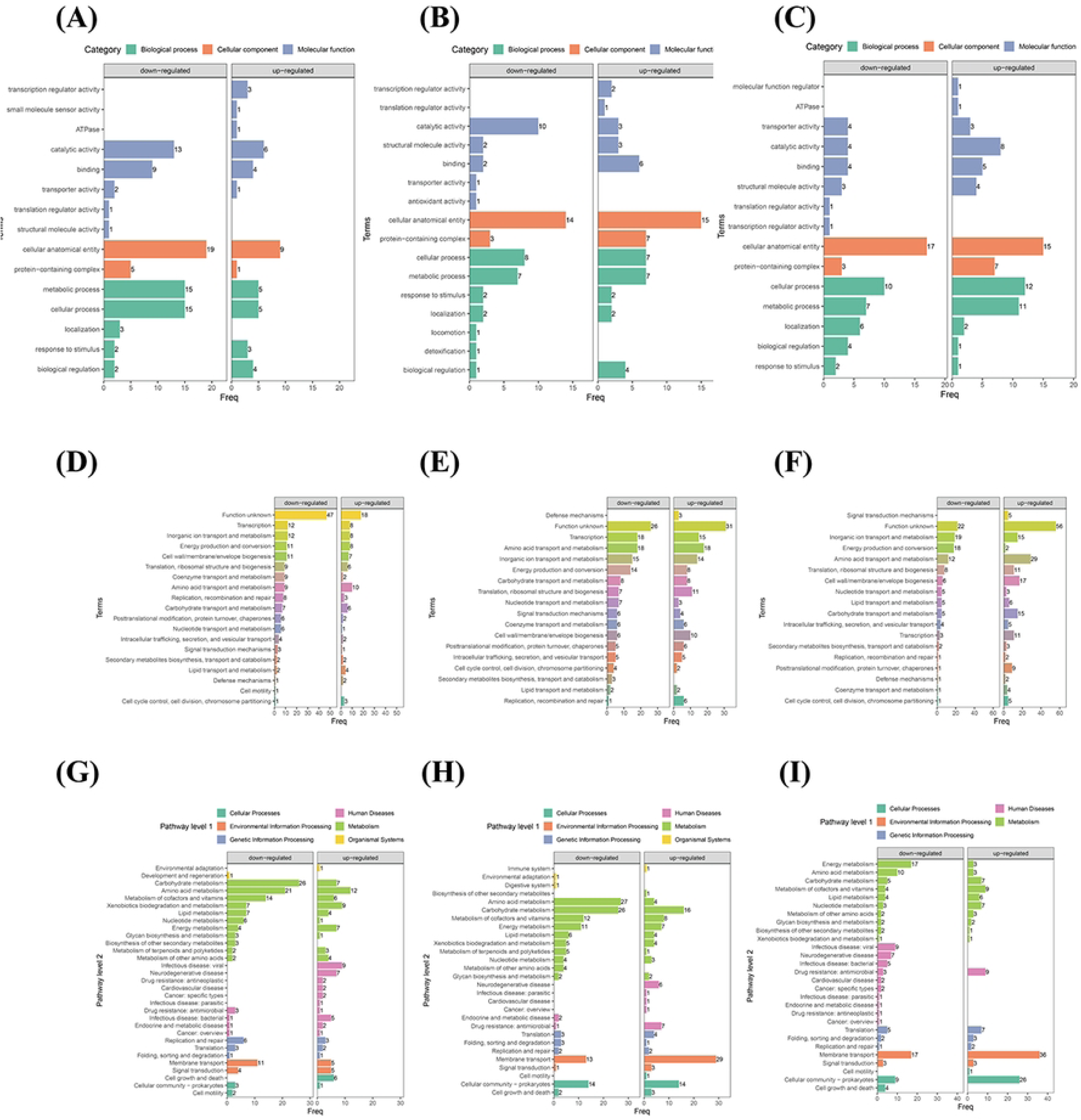
(A) Gene Ontology (GO) functional classification of differentially expressed prot eins in the 80°C/20 min vs. 95°C/20 min comparative group. (B) Gene Ontology (GO) fu nctional classification of differentially expressed proteins in the 0.6% HCHO/48 h vs. 0.6 % HCHO/72 h comparative group. (C) Gene Ontology (GO) functional classification of di fferentially expressed proteins in the heat-inactivated vs. formaldehyde-inactivated comparati ve group. (D) KOG functional classification of differentially expressed proteins in the 80° C/20 min vs. 95°C/20 min comparative group. (E) KOG functional classification of differe ntially expressed proteins in the 0.6% HCHO/48 h vs. 0.6% HCHO/72 h comparative group. (F) KOG functional classification of differentially expressed proteins in the heat-inactiva ted vs. formaldehyde-inactivated comparative group. (G) KEGG functional classification of differentially expressed proteins in the 80°C/20 min vs. 95°C/20 min comparative group. (H) KEGG functional classification of differentially expressed proteins in the 0.6% HCHO/48 h vs. 0.6% HCHO/72 h comparative group. (I) KEGG functional classification of differ entially expressed proteins in the heat-inactivated vs. formaldehyde-inactivated comparative group.

For the 0.6% formaldehyde treatment, proteins with higher levels in the 48 h group compared to the 72 h group were primarily enriched in binding functions (6 genes) and transporter activity (4 genes). In contrast, proteins with lower levels in the 0.6% formaldehyde/48 h group relative to the 0.6% formaldehyde/72 h group were concentrated in catalytic activity (10 genes) and cellular anatomical entity pathways (14 genes) (Figure 3B).

Proteins exhibiting higher expression in the heat-inactivated group compared to the formaldehyde-treated group were highly enriched in cellular anatomical entities (17 genes) and cellular processes (10 genes). This suggests that heat inactivation may be more conducive to maintaining or promoting the compensatory expression of proteins related to cellular structure and function, whereas formaldehyde treatment might lead to the downregulation or functional impairment of these proteins. Conversely, proteins with lower expression in the heat-inactivated group relative to the formaldehyde-treated group were significantly enriched in catalytic activity (8 genes), cellular processes (12 genes), metabolic processes (11 genes), and cellular anatomical entities (15 genes) (Figure 3C).

KOG functional classification analysis indicated that proteins at higher levels in the 80°C/20 min group compared to the 95°C/20 min group were predominantly associated with “amino acid transport and metabolism” (10 genes) and “lipid transport and metabolism” (4 genes). In contrast, proteins with lower expression in the 80°C/20 min group relative to the 95°C/20 min group were significantly enriched in transcription, inorganic ion transport and metabolism, energy production and conversion, and cell wall/membrane/envelope biogenesis pathways (Figure 3D).

Under 0.6% formaldehyde treatment, comparison between the 48 h and 72 h groups revealed that 31 proteins with higher levels in the 48 h group had unknown functions, while others were primarily concentrated in pathways such as transcription, amino acid transport and metabolism, and inorganic ion transport and metabolism. Similarly, proteins with lower levels in the 48 h group relative to the 72 h group included 26 proteins of unknown function, with others primarily involved in transcription, amino acid transport and metabolism, and inorganic ion transport and metabolism pathways (Figure 3E).

Proteins with higher levels in the heat-inactivated group compared to the formaldehyde-treated group were enriched in inorganic ion transport and metabolism and energy production and conversion pathways. Conversely, proteins with lower levels in the heat-inactivated group relative to the formaldehyde-treated group were primarily concentrated in amino acid transport and metabolism and cell wall/membrane/envelope biogenesis pathways (Figure 3F).

KEGG pathway classification analysis showed that proteins with higher levels in the 80°C/20 min group relative to the 95°C/20 min group were enriched in amino acid metabolism (12 genes) and lipid metabolism (9 genes pathways). Conversely, proteins with lower levels in the 80°C/20 min group compared to the 95°C/20 min group were significantly enriched in carbohydrate metabolism (26 genes), amino acid metabolism (21 genes), and membrane transport (11 genes) pathways (Figure 3G).

Under 0.6% formaldehyde treatment, comparison between the 48 h and 72 h groups indicated that proteins with relatively higher levels in the 48 h group were significantly enriched in membrane transport (29 genes) and cellular communities—prokaryotes (14 genes) pathways. Proteins with relatively lower levels in the 48 h group were enriched in amino acid metabolism (27 genes), carbohydrate metabolism (26 genes), and energy metabolism (11 genes) pathways (Figure 3H).

Proteins with higher levels in the heat-inactivated group relative to the formaldehyde-treated group were highly active in membrane transport (36 genes) and cellular communities (26 genes) pathways. Conversely, proteins with lower levels in the heat-inactivated group compared to the formaldehyde-treated group were enriched in energy metabolism (17 genes) and membrane transport (17 genes) pathways (Figure 3I).

### 2.5 Prediction and Classification Based on Protein Subcellular Localization

Subcellular localization analysis was performed to elucidate the spatial distribution characteristics of differentially expressed *Brucella* proteins under various inactivation conditions. It was found that proteins exhibiting higher levels in the 80°C/20 min group, as compared to the 95°C/20 min group, were relatively enriched in plasma membrane-associated proteins (5 genes). Conversely, proteins with lower levels in the 80°C/20 min group, relative to the 95°C/20 min group, were significantly enriched in the cytoplasm (13 genes) and plasma membrane (3 genes). This suggests that a “partially denatured-partially aggregated” mixed state may be formed at 80°C, while a uniformly and completely denatured state is likely achieved at 95°C (Figure 4A).

**Figure 4:**
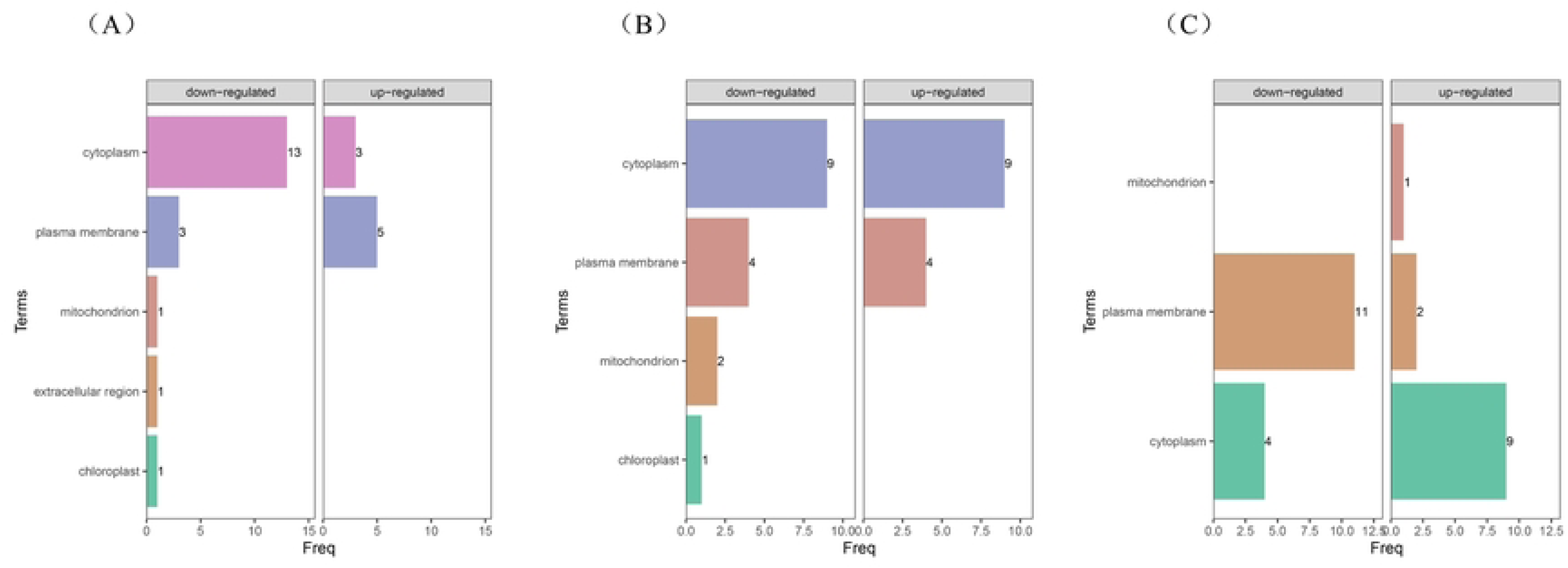
(A) Subcellular localization analysis of differentially expressed proteins in the 8 0°C/20 min vs. 95°C/20 min comparative group. (B) Subcellular localization analysis of di fferentially expressed proteins in the 0.6% HCHO/48 h vs. 0.6% HCHO/72 h comparative group. (C) Subcellular localization analysis of differentially expressed proteins in the heat-inactivated vs. formaldehyde-inactivated comparative group.

Under 0.6% formaldehyde treatment, a comparison between the 48 h and 72 h groups revealed that proteins at relatively higher levels in the 48 h group primarily belonged to cytoplasmic proteins (9 genes). Conversely, proteins at relatively lower levels were predominantly found in the cytoplasm (9 genes) and mitochondria (2 genes). This indicates that the 48 h treatment primarily resulted in cytoplasmic soluble phases, whereas signals appeared at 72 h as a result of subcellular organelle disintegration (Figure 4B).

Only a small number of proteins (2 genes) with higher levels in the heat-inactivated group, relative to the formaldehyde-treated group, were distributed in the plasma membrane. In contrast, proteins with lower levels in the heat-inactivated group, compared to the formaldehyde-treated group, were concentrated in the plasma membrane (11 genes) and cytoplasm (9 genes). This suggests that heat inactivation preferentially enhances the detection of a few soluble membrane proteins but leads to signal attenuation for most membrane and cytoplasmic proteins due to aggregation. Conversely, formaldehyde inactivation preserves a broader range of epitopes by stabilizing proteins in a soluble state (Figure 4C).

### 2.6 Functional enrichment analysis of differentially expressed proteins

Biological process enrichment analysis revealed that proteins with lower levels in the 80°C/20 min group, compared to the 95°C/20 min group, exhibited limited enrichment in the biological processes studied. In contrast, proteins with higher levels in the 80°C/20 min group, relative to the 95°C/20 min group, were significantly enriched in processes such as the regulation of cellular macromolecule biosynthetic processes and the regulation of gene expression (Figure 5A).

**Figure 5:**
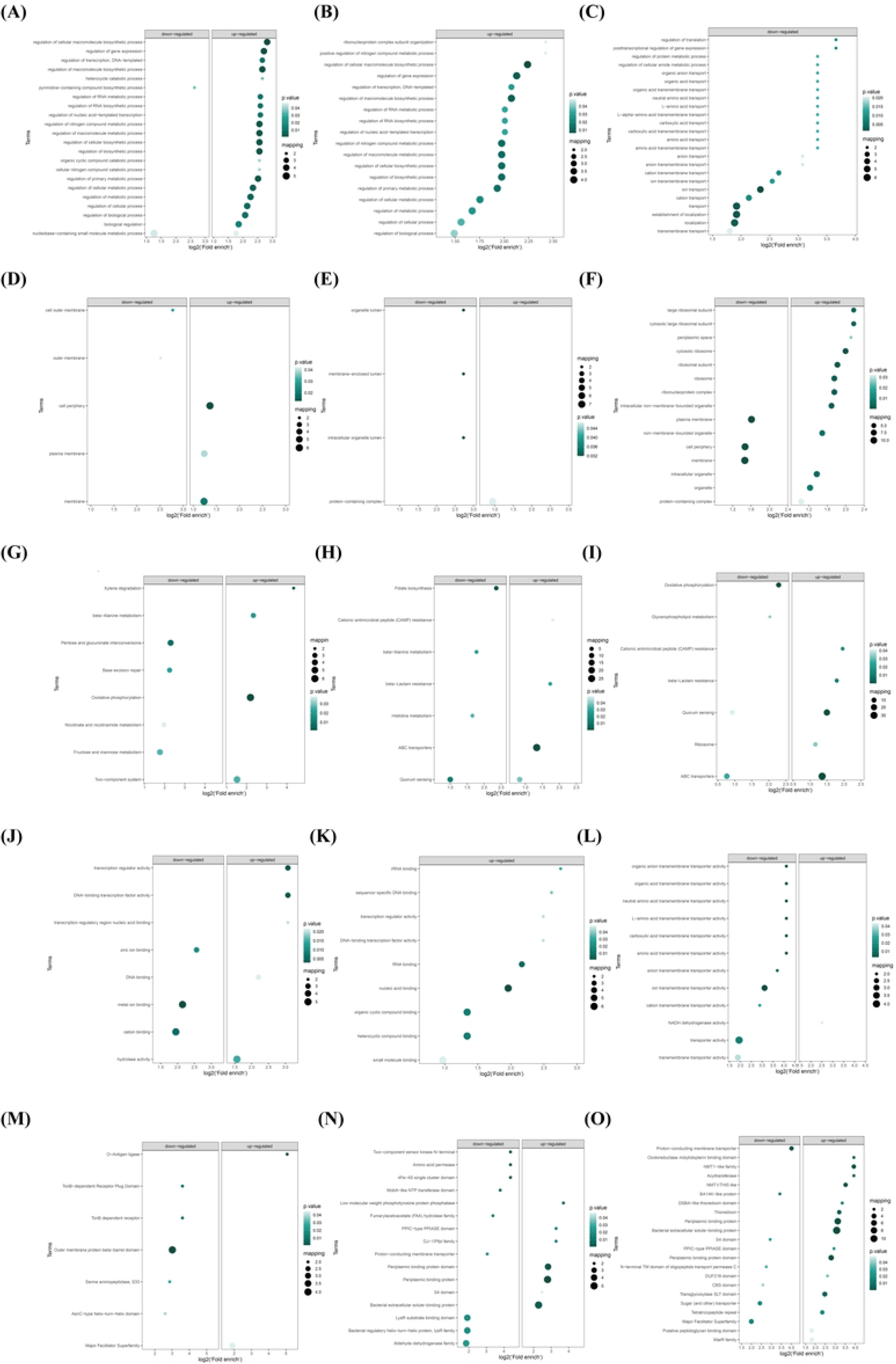
(A) Biological process enrichment of differentially expressed proteins in the 80° C/20 min vs. 95°C/20 min comparative group. (B) Biological process enrichment of differ entially expressed proteins in the 0.6% HCHO/48 h vs. 0.6% HCHO/72 h comparative gro up. (C) Biological process enrichment of differentially expressed proteins in the heat-inacti vated vs. formaldehyde-inactivated comparative group. (D) Cellular component enrichment of differentially expressed proteins in the 80°C/20 min vs. 95°C/20 min comparative group. (E) Cellular component enrichment of differentially expressed proteins in the 0.6% HCH O/48 h vs. 0.6% HCHO/72 h comparative group. (F) Cellular component enrichment of di fferentially expressed proteins in the heat-inactivated vs. formaldehyde-inactivated comparati ve group. (G) Molecular function enrichment of differentially expressed proteins in the 80° C/20 min vs. 95°C/20 min comparative group. (H) Molecular function enrichment of differ entially expressed proteins in the 0.6% HCHO/48 h vs. 0.6% HCHO/72 h comparative gro up. (I) Molecular function enrichment of differentially expressed proteins in the heat-inacti vated vs. formaldehyde-inactivated comparative group. (J) KEGG pathway enrichment of differentially expressed proteins in the 80°C/20 min vs. 95°C/20 min comparative group. (K) KEGG pathway enrichment of differentially expressed proteins in the 0.6% HCHO/48 h vs. 0.6% HCHO/72 h comparative group. (L) KEGG pathway enrichment of differentially expressed proteins in the heat-inactivated vs. formaldehyde-inactivated comparative group. (M) Pfam domain enrichment of differentially expressed proteins in the 80°C/20 min vs. 9 5°C/20 min comparative group. (N) Pfam domain enrichment of differentially expressed pr oteins in the 0.6% HCHO/48 h vs. 0.6% HCHO/72 h comparative group. (O) Pfam domai n enrichment of differentially expressed proteins in the heat-inactivated vs. formaldehyde-in activated comparative group.

Under 0.6% formaldehyde treatment, comparison between the 48 h and 72 h groups indicated that multiple regulation-related biological processes, including gene expression regulation, transcription regulation, and RNA metabolic process regulation, were significantly enriched among proteins at relatively higher levels in the 48 h group. No significant enrichment was observed for proteins at relatively lower levels (Figure 5B).

Proteins with lower levels in the heat-inactivated group, relative to the formaldehyde-treated group, were significantly enriched in biological processes such as ion transport, cation transport, and transmembrane transport. Conversely, no enrichment in biological processes was found for proteins with higher levels in the heat-inactivated group compared to the formaldehyde-treated group (Figure 5C).

Through Cellular Component enrichment analysis, it was observed that proteins with lower levels in the 80°C/20 min group, compared to the 95°C/20 min group, showed relatively low significance and fewer mapped proteins in their enrichment, with only a limited degree of enrichment in the cell outer membrane and outer membrane. Overall enrichment and significance were modest. Conversely, proteins with higher levels in the 80°C/20 min group, relative to the 95°C/20 min group, exhibited significant enrichment in cellular components such as the cell periphery, plasma membrane, and membrane (Figure 5D).

Under 0.6% formaldehyde treatment, a comparison between the 48 h and 72 h groups revealed that proteins with relatively lower expression at 48 h were primarily enriched in membrane-enclosed lumens, organelle lumens, and intracellular organelle lumens. In contrast, proteins with relatively higher expression at 48 h were predominantly enriched in protein-containing complex-related regions (Figure 5E).

Proteins with lower levels in the heat-inactivated group, relative to the formaldehyde-treated group, were significantly enriched in membrane-associated components, including the plasma membrane, membrane, and cell periphery. Proteins with higher levels in the heat-inactivated group were significantly enriched in ribosome and protein complex-related components, such as the large ribosomal subunit, cytosolic ribosome, ribonucleoprotein complex, and protein-containing complex (Figure 5F).

Molecular function enrichment analysis indicated that proteins with lower levels in the 80°C/20 min group, compared to the 95°C/20 min group, were enriched in metal ion binding-related molecular functions, specifically zinc ion binding, metal ion binding, and cation binding. Conversely, proteins with higher levels in the 80°C/20 min group, relative to the 95°C/20 min group, were predominantly enriched in pathways related to transcriptional regulation activity, such as transcription regulator activity, DNA-binding transcription factor activity, and transcription regulatory region nucleic acid binding, as well as DNA binding-related hydrolase activity pathways (Figure 5G).

Under 0.6% formaldehyde treatment, comparison between the 48 h and 72 h groups revealed that multiple regulation-related molecular functions were significantly enriched among proteins at relatively higher levels in the 48 h group. These included rRNA binding, sequence-specific DNA binding, transcription regulator activity, DNA-binding transcription factor activity, RNA binding, nucleic acid binding, organic cyclic compound binding, heterocyclic compound binding, and small molecule binding. No significant enrichment was observed for proteins at relatively lower levels (Figure 5H).

Proteins with lower levels in the heat-inactivated group, relative to the formaldehyde-treated group, were significantly enriched in various types of transmembrane transport, including organic anion transmembrane transport, organic acid transmembrane transport, neutral amino acid and L-amino acid transmembrane transport, and carboxylic acid transmembrane transport. Minimal significant enrichment was found for molecular functions of proteins at relatively higher levels in the heat-inactivated group, with only a slight enrichment observed in the NADH dehydrogenase activity pathway (Figure 5I).

KEGG pathway enrichment analysis revealed that proteins with lower levels in the 80°C/20 min group, compared to the 95°C/20 min group, were enriched in pathways such as Xylene degradation, beta-Alanine metabolism, Pentose and glucuronate interconversions, Base excision repair, Oxidative phosphorylation, Nicotinate and nicotinamide metabolism, Fructose and mannose metabolism, and Two-component system. Conversely, proteins with higher levels in the 80°C/20 min group, relative to the 95°C/20 min group, were primarily enriched in Oxidative phosphorylation, beta-Alanine metabolism, and Xylene degradation (Figure 5J).

Under 0.6% formaldehyde treatment, comparison between the 48 h and 72 h groups indicated that proteins at relatively lower levels in the 48 h group were enriched in Folate biosynthesis, Beta-Alanine metabolism, Histidine metabolism, and Quorum sensing pathways. Proteins at relatively higher levels in the 48 h group were enriched in ABC transporters, Beta-Lactam resistance, and Cationic antimicrobial peptide (CAMP) resistance pathways (Figure 5K).

Proteins with lower levels in the heat-inactivated group, relative to the formaldehyde-treated group, were significantly enriched in Oxidative phosphorylation, Glycerophospholipid metabolism, Quorum sensing, and ABC transporters pathways. Proteins with higher levels in the heat-inactivated group, compared to the formaldehyde-treated group, were enriched in Beta-Lactam resistance, Cationic antimicrobial peptide (CAMP) resistance, and Ribosome pathways (Figure 5L).

Pfam domain enrichment analysis revealed that proteins with lower levels in the 80°C/20 min group, compared to the 95°C/20 min group, exhibited low enrichment in domains such as O-Antigen ligase, TonB-dependent Receptor Plug Domain/TonB dependent receptor, Outer membrane protein beta-barrel domain, Serine aminopeptidase, AsnC-type helix-turn-helix domain, and Major Facilitator Superfamily. In contrast, only the O-Antigen ligase family was enriched among proteins with higher levels in the 80°C/20 min group (Figure 5M).

Under 0.6% formaldehyde treatment, comparison between the 48 h and 72 h groups indicated that proteins at relatively higher levels in the 48 h group included domains such as Bacterial extracellular solute-binding protein, Periplasmic binding protein domain/Periplasmic binding protein, LysR substrate binding domain & LysR family regulatory protein, Aldehyde dehydrogenase family, and S4 domain. Conversely, proteins at relatively lower levels were significantly enriched in Two-component sensor kinase N-terminal, Amino acid permease, Iron-sulfur cluster domain (4Fe-4S single cluster domain), MobA-like NTP transferase domain, FAA hydrolase family, DJ-1/PfpI family, and PPIASE domain, among others (Figure 5N).

The Pfam functions enriched in proteins with lower levels in the heat-inactivated group, relative to the formaldehyde-treated group, primarily involved transport-related domains/families, electron transport and redox-related functions, peptidoglycan binding and cell wall-related domains, and transport/binding protein domains. Proteins with higher levels in the heat-inactivated group, compared to the formaldehyde-treated group, were enriched in thioredoxin-related domains, peripheral membrane binding protein-related domains, and methyltransferase-related domains (Figure 5O).

### 2.7 Protein–protein interaction (PPI) networks

Investigating protein-protein interaction (PPI) regulatory networks from a bioinformatics perspective facilitates a deeper understanding of the molecular mechanisms underlying the effects of different inactivation methods on *Brucella* proteins. Within PPI networks, highly clustered proteins often exhibit similar functions and perform biological roles through synergistic interactions.

In the protein-protein interaction (PPI) network derived from the 80°C/20 min inactivation group and the 95°C/20 min inactivation group, two main modules were identified: a ribosome-related protein module and an NADH dehydrogenase complex-related protein module. The ribosome-related protein module primarily contained high-level expression nodes such as rpmE, rpmF, hflX, and rsfS. Conversely, low-level expression nodes included rpmC, infC, rpmD, and rpmI. These proteins are largely ribosomal subunit proteins involved in protein synthesis. The NADH dehydrogenase complex-related protein module featured key high-level expression nodes like nuoM, nuoA, nuoH, cydA, ccoN, cycA, and BAB1_0389, while major low-level expression nodes included BAB1_0833, BAB2_0733, and BAB2_0837. These proteins constitute respiratory chain Complex I (NADH dehydrogenase) and cytochrome complexes, which are crucial for energy metabolism (Figure 6A).

**Figure 6:**
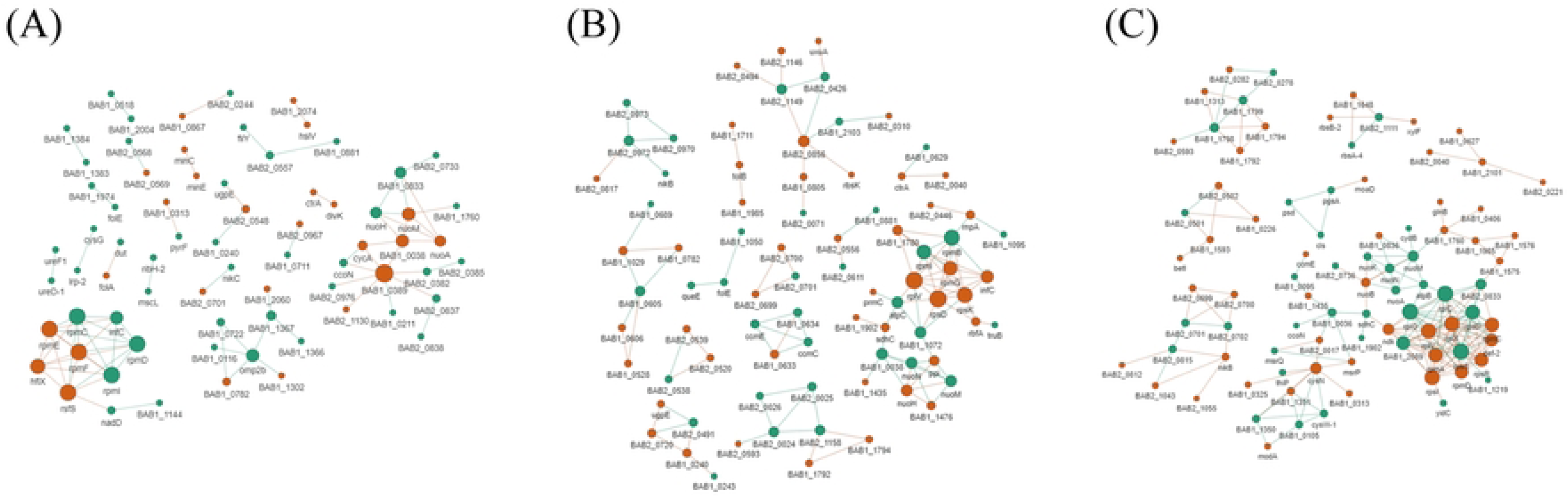
(A) Protein-protein interaction network of the 80°C/20 min vs. 95°C/20 min co mparative group. (B) Protein-protein interaction network of the 0.6% HCHO/48 h vs. 0.6% HCHO/72 h comparative group. (C) Protein-protein interaction network of the heat-inactiv ated vs. formaldehyde-inactivated comparative group.

Within the protein-protein interaction (PPI) network comparing the 0.6% formaldehyde inactivation 48 h group and the 0.6% formaldehyde inactivation 72 h group, two prominent clusters were identified: a ribosome-related cluster (rpmG, rpmJ, rplV, rpsD, rpsK) and an energy metabolism-related protein cluster (atpC; sdhC; nuoH, nuoM, nuoN). The ribosome-related proteins exhibited a large aggregation of red nodes (high level), including rpmG, rpmI, rpsK, rpsD, infC, rnl, rnpA, and rnpB, all of which are involved in protein synthesis. Their high interconnectivity and centrality suggest a generally high level of ribosomal proteins at 48 h. The energy metabolism-related protein subpopulation, comprising proteins such as nuoH, nuoM, nuoN, sdhC, and atpC, which are associated with the respiratory chain and energy metabolism, were predominantly green in the network, indicating lower protein levels after 48 h of inactivation (Figure 6B).

In the PPI network of differentially expressed proteins from the heat inactivation and formaldehyde inactivation groups, ribosomal protein clusters (rpl, rps, rpm) exhibited a mixed distribution of high and low expression, indicating significant differences in ribosomal protein expression under heat inactivation conditions (Figure 6C). The respiratory chain and metabolism-related protein cluster, which includes proteins such as nuo, sdhC, and ccmE involved in energy metabolism and respiratory chain assembly, displayed a higher number of low-expression nodes. This suggests that heat inactivation, compared to formaldehyde inactivation, more readily leads to the destruction or degradation of cell energy metabolism-related proteins (Figure 6C). The transport and response protein cluster contained numerous proteins associated with substance transport and metabolic regulation (nikB, betI, modA), showing a mixed high and low expression pattern, which reflects complex regulation of these functional proteins between the two groups (Figure 6C). Interestingly, modA was found at a relatively high level, suggesting a protective effect of heat inactivation on certain specific transport proteins, thereby favoring the retention of binding and transport functions for some metal ions or nutrients.

## 3 Discussion

Despite its significance as a zoonotic pathogen, scientific research and investigations concerning *Brucella* have progressed relatively slowly over the years. A notable impediment to this research is the requirement for Biosafety Level 3 (BSL-3) containment for handling pathogenic *Brucella* [21], which significantly restricts experimental procedures. Furthermore, *Brucella* has not garnered a primary research focus among bacterial pathogens commensurate with its pathogenic impact. In studies addressing crucial aspects of *Brucella* pathogenesis and immune evasion, inactivation of the bacterium is a frequently encountered preliminary step, particularly during in vitro validation experiments. It has been observed that the “Diagnosis and Treatment Protocol for Brucellosis (2023 Edition)” issued by China’s National Health Commission states that *Brucella* can be inactivated by moist heat at 60°C for 10-20 minutes. However, previous studies on the heat inactivation of cultured *Brucella* indicated that boiling temperatures for over an hour, or the use of strong acids such as CHCA, were required for complete inactivation [22]. This significant discrepancy in temperature and time requirements not only creates ambiguity regarding laboratory safety and sample inactivation procedures for *Brucella* but also highlights the complexity of its inactivation parameters. Such differences may arise from various factors, including the research subject (pure cultures versus clinical or environmental samples), bacterial strain type, heating medium, and the methods used to assess inactivation efficacy. It can be inferred that different inactivation methods will inevitably exert distinct effects on samples. To provide more precise guidance for *Brucella* laboratory safety, vaccine development, and disease control practices, a systematic and in-depth investigation into the survival characteristics of *Brucella* under diverse inactivation conditions is deemed necessary. In this study, the biosafety of four inactivation methods was initially evaluated, leading to the identification of methods capable of inactivating the Rev.1 vaccine strain. Subsequently, analysis of clinical isolates revealed incomplete inactivation in the 0.4% HCHO for 72 h group. This incomplete inactivation may be attributed to the enhanced tolerance of clinical wild-type strains, which possess superior compensatory transport capabilities, biofilm protection, stress protein expression, and quorum sensing regulation. In contrast, the Rev.1 vaccine strain, due to prolonged artificial attenuation, exhibits weaker resistance mechanisms, rendering it more susceptible to low concentrations of formaldehyde [23]. This finding underscores that inactivation parameters established for vaccine strains or a single strain cannot be directly extrapolated to other clinical samples; specific analysis for each target strain may be required to determine unique parameters. While high-quality samples are essential for research, based on our analysis, for inactivation of *Brucella* in grassroots medical settings or clinical diagnoses, the use of more rigorous methods is recommended, such as sodium hypochlorite, sodium hydroxide, phenols, in combination with ultraviolet irradiation. If formaldehyde is to be employed for inactivation, higher concentrations and extended exposure times (up to several hours) are advised [24].

As a pathogenic organism posing a significant biosafety threat, the manipulation and study of *Brucella* necessitate Biosafety Level 3 (BSL-3) laboratory operations and stringent physical containment measures. Such facilities are often not readily available in basic veterinary units within countries burdened by high *Brucella* prevalence, particularly in the Global South [25]. Consequently, the development of a reliable *Brucella* inactivation method is expected to enhance biosafety for laboratory personnel and facilitate *Brucella* research [26]. Through a robust and straightforward inactivation process, samples can be safely rendered non-infectious prior to being transported to laboratories equipped with high-end instrumentation, such as mass spectrometers, thereby enabling further investigation [27]. Previous research has focused on the development of the CHCA method for direct mass spectrometric identification of *Brucella*; however, the impact of this method on antigenicity was not described. In the current study, following initial screening by ELISA and Western blot, it was observed that both heat inactivation and formaldehyde inactivation methods preserved the reactivity of *Brucella* proteins to a significantly greater extent than the CHCA method. Therefore, antigens inactivated by these methods were subsequently utilized for immunoprecipitation (IP) followed by mass spectrometric analysis.

Immunoprecipitation-mass spectrometry has been established as a powerful tool for identifying protein-protein interactions, while emerging immunoproteomics significantly contributes to advancing vaccine development [28]. Although previous mass spectrometry methods had certain limitations, the recent advent of Astral-DIA has revitalized proteomics research due to its exceptional stability and sensitivity [29]. To systematically evaluate the impact of different *Brucella* inactivation methods on bacterial antigenicity and proteomic integrity, a detailed analysis was conducted using antibody immunoprecipitation coupled with mass spectrometry from an Astral-DIA proteomic perspective. Following mass spectrometric analysis, it was determined that approximately 60% protein coverage (approximately 2000 out of 3300 proteins) was achieved with Astral-DIA. This coverage is comparable to, or even surpasses, that typically obtained from direct mass spectrometric analysis of whole-cell lysate proteins [30].

In the heat inactivation groups (80°C/95°C), significant protein denaturation and aggregation were observed. Proteomics analysis, including PPI network analysis, revealed substantial differences in the expression of ribosomal proteins (e.g., rpmE, rpmF) and energy metabolism proteins (e.g., nuoM, sdhC) within these groups. While catalytic activity and metabolic process-related proteins were partially retained in the 80°C group, a more uniform denaturation state was evident in the 95°C group, accompanied by a diminished signal from plasma membrane proteins. This suggests that the detrimental effects of heat inactivation on conformational epitopes may be attributed to the disruption of hydrogen bond networks at high temperatures, leading to the aggregation of key antigens (e.g., outer membrane proteins OMP25/31) and a reduction in their immunoreactivity [31]. KEGG enrichment analysis further indicated an upregulation of the Beta-Lactam resistance pathway, but a downregulation of Oxidative phosphorylation and Quorum sensing pathways, implying impaired energy metabolism and virulence factor function. Additionally, an upregulation of inorganic ion transporters (e.g., zntA zinc transporter) was detected in the heat inactivation groups, co-expressed with hflX (heat shock GTPase) in the PPI network. This may suggest a link between metal ion homeostasis and stress response, potentially offering new targets for enhancing the efficacy of heat inactivation through metal chelators, which will be investigated in future studies [32].

In the formaldehyde-inactivated groups (0.6% for 48h/72h), it was observed that formaldehyde stabilized protein structures through amino cross-linking, thereby better preserving soluble antigens [33]. Differential expression analysis revealed that ribosomal proteins (rpmG, rpsK) and regulatory proteins (e.g., transcription factors) were highly expressed in the 48h group, with subcellular localization predominantly in the cytoplasmic soluble phase, which is conducive to epitope exposure. Among the enriched molecular functions, DNA binding and RNA binding pathways were significantly upregulated, indicating superior preservation of nucleic acid-related antigens by formaldehyde. However, extending the inactivation period to 72h led to the disintegration of subcellular organelles, resulting in the release of mitochondrial protein signals, which could introduce non-specific background noise. Although formaldehyde inactivation demonstrated better antigen preservation, residual toxicity (e.g., unreacted formaldehyde) may necessitate stringent neutralization treatment to prevent interference with immunological experiments [34]. A comprehensive comparison indicated that formaldehyde inactivation outperformed heat inactivation in terms of antigen reactivity. The heat inactivation group exhibited 318 differentially expressed proteins compared to the formaldehyde group. Proteins upregulated by heat inactivation were predominantly enriched in cellular structure and function (e.g., cellular anatomical entity), whereas downregulated proteins involved critical transport and metabolic pathways (e.g., ABC transporters), reflecting irreversible damage to antigenic epitopes. Conversely, formaldehyde inactivation, by maintaining proteins in a soluble state through mild cross-linking, may be more suitable for immunology-related studies requiring antigenicity preservation [35].

Concurrently, it was observed that several proteins with previously uncharacterized functions exhibited significantly differential expression levels. These “dark matter proteins” are hypothesized to represent a novel reservoir of antigens. Subsequent investigations will focus on identifying their epitopes and functions through biological and immunological assays.

Despite experimental data indicating a clear disadvantage for heat-inactivated groups (80°C/95°C for 20 min) compared to formaldehyde-inactivated groups (0.6% for 48h/72h) in terms of protein integrity and epitope preservation [36], the distinct PPI profiles associated with different inactivation methods suggest their complementary application potential. The unique value of heat inactivation lies in its ability to degrade energy metabolism and cytoplasmic proteins while specifically preserving and enhancing the immunogenicity of certain membrane proteins (e.g., virulence factor ModA) and stress response proteins (e.g., heat-induced ribosomal proteins) [37]. These proteins could serve as “danger signals” to activate innate immunity or as targeted antigens, making them relevant for the development of subunit vaccines or for studies exploring the adjuvant effects of heat shock proteins [38]. Conversely, the primary advantage of formaldehyde inactivation is its capacity to maintain protein solubility and the integrity of multifunctional antigenic epitopes. Specifically, cross-linking stabilizes epitope-rich ribosomal protein-nucleic acid complexes and cytoplasmic soluble proteins, making it suitable for research on whole-cell inactivated vaccines that require conformational epitope preservation or for multi-epitope antigen screening [39]. In summary, heat inactivation primarily focuses on exposing and enhancing the immune reactivity of specific “danger signals” or virulence factors, whereas formaldehyde inactivation excels at preserving a broad range of multi-epitopic protein structural integrity. This highlights that the choice of inactivation method should be guided by the priorities of the downstream application (e.g., antigen breadth vs. specific epitopes, safety vs. operational complexity), rather than by a single performance metric. Further experimental validation of the actual mechanisms of differentially expressed proteins is warranted to optimize precise prevention and control strategies for brucellosis.

## 4. Conclusion

In summary, this study systematically investigated the impact of *Brucella* inactivation processes on biosafety and antigenic reactivity from an antibody-immunoprecipitation-mass-spectrometry-based Astral-DIA proteomic perspective. It was determined that inactivation parameters must be precisely matched to strain characteristics, and parameters derived from vaccine strains cannot be directly extrapolated. Furthermore, antigen preservation was found to be method-dependent, with distinct differences observed in the preservation biases of various methods. The advantages of Astral-DIA technology in validating immunoproteomics were re-affirmed, and for the first time, the dynamic analysis of protein-protein interaction networks post-inactivation was achieved, providing molecular evidence for the mechanisms of antigenicity loss. However, the current PPI network analysis has not yet encompassed the dynamic damage processes of critical conformational epitopes; therefore, further biological experimental techniques may be required for a more comprehensive assessment in the future. These findings will be disseminated to grassroots laboratories and integrated into *Brucella* mechanistic research through more rigorous experimental studies, with the aim of providing precise guidance for *Brucella* vaccine development and quality control, thereby establishing a solid scientific basis for more effective disease prevention and control strategies.

## Ethics Statement

The authors have nothing to report.

## Consent

All authors approved the manuscript.

## Acknowledgments

This work was supported by grants from the Key Research and Development Program of Gansu Province (25YFNA015) and the Agricultural Science and Technology Support Project of the Department of Agriculture and Rural Affairs of Gansu Province (KJZC-2024-13) and the Innovation Program of Chinese Academy of Agricultural Sciences (CAAS-ZDRW-202410) and the Major Science and Technology Project of Gansu Province (23ZDNA007).

## Author contributions

Yuefeng Chu and Fuying Zheng designed the study. Yijian Liu, Jiazhen Ge and Guodong Song conducted the experiments. Yijian Liu and Jiazhen Ge draft the manuscript. Pengcheng Gao isolated the strain, and Mengzhu Qi undertook the serum collection. Wenhao Wang, Yingying Xie, Ziqing Wang and Renge Li contributed to optimizing the experimental conditions.

